# Differential effects of ischemia and inflammation on plasma-derived extracellular vesicle characteristics and function

**DOI:** 10.1101/2025.05.13.653696

**Authors:** Yvonne Couch

**Author notes:** Correspondence addressed to: **Associate Professor Yvonne Couch**, Oxford BHF Centre of Research Excellence Transitional Research Fellow, Nuffield Department of Clinical Neurosciences, University of Oxford, Dorothy Crowfoot-Hodgkin Building, Sherrington Road, Oxford, OX1 3QU, UK.

## Abstract

Extracellular vesicles (EVs) have long been understood to be important mediators of cell-to-cell communication and may lead to the molecular aftermath and exacerbation of brain injuries such stroke. This study explored how the source of the EVs influenced their characteristics and the effect these differences had on naïve brain tissue. EVs were isolated from animals post-stroke in the acute or chronic stages of recovery in animals with and without reperfusion, and from a model of systemic inflammation (i.p. lipopolysaccharide). The data show that neither stroke nor inflammation significantly increase EV numbers compared to sham or naïve animals. Post-stroke EVs exhibited a panel of different platelet and inflammatory markers, when compared to EVs derived from a model of inflammation, reflecting differences between stroke and systemic immune activation. When injected into the brain, both stroke-derived and inflammation-derived EVs induced expression of pro-inflammatory cytokine gene expression, suggesting a potential role in neuroinflammation. However, there was a lack of distinct glial and astrocyte reactivity in response to any EVs, despite robust changes in ICAM reactivity. The findings here underscore the complexity of EV roles in pathophysiology and highlight the need for improved EV isolation methods. With further longitudinal studies we may be able to more accurately determine how the context of the injury (reperfusion vs no reperfusion vs inflammation) might contribute to the EV populations and their function. Understanding more about EVs in different contexts will improve our ability to use EVs as biomarkers, but also our capacity to interfere with EV biology as a novel therapeutic approach.

## Introduction

Ischemia within the central nervous system (CNS) results in the rapid death of neurons and the activation of other key cells within the neurovascular unit and brain parenchyma. These include not only microglia and astrocytes, but also cells such as endothelia and pericytes. Pathologically, cerebral ischemia has significant long-term consequences for brain health. Studies of the inflammatory status of the brain in the chronic phases post-stroke have suggested that activation of microglia and astrocytes persists far beyond what might be considered the acute ‘clean up’ phase after injury (1, 2). This chronic CNS inflammation is likely to contribute to the development of neurodegeneration and cognitive decline.

In neurodegenerative diseases such as Alzheimer’s and Parkinson’s, chronic inflammation is thought to contribute to the proteinaceous nature of the pathology (3, 4). In vascular dementia, where there is chronic vascular pathology within the CNS, inflammation may also play a crucial role (5, 6). In stroke, we have a vascular event which has the capacity to precipitate inflammation and long-term vascular changes but the nature of how this event is communicated throughout the brain is only just beginning to be understood (7, 8).

Extracellular vesicles (EVs) are lipid bound vesicles which are an important novel means of cell-to-cell communication (9, 10). They have traditionally been categorized according to their biogenesis, with – for example - ‘exosomes’ released from multivesicular bodies and ‘microvesicles’ blebbed from the cell surface (11). However, the inherent challenges associated with isolating these individual populations of vesicles has necessitated the use of the term extracellular vesicle to cover the majority of the subtypes currently understood (11, 12). EVs have been known to be functional since the late 1980s and more recently they have been found to contribute to the exacerbation of inflammation after acute CNS injuries (13–16). This promising preliminary data suggests that EVs could act as a signature for the injury and could be used as biomarkers for long-term outcomes (17). Furthermore, the injury signature is likely to dictate the downstream function of the EV. However, because the study of EVs in CNS diseases is still a relatively new field, a large number of the published studies to date have used differential centrifugation to isolate EV populations, something which is known to increase protein contamination (12, 18). Here, we wished to determine the degree to which a ‘purer’ population of particles contributed to CNS inflammation, and the degree to which the precipitating factors in the release of those particles, i.e. the nature of the injury, contributed to their functional effects. Understanding these aspects of EV biology will enable us to make more effective decisions on the use of EVs as biomarkers (17), and on the function of EVs in cell-to-cell communication.

## Materials and Methods

### Animals

Male CD1 mice, 8–10 weeks of age, were housed under standard diurnal lighting conditions (12 h) with *ad libitum* access to food and water. All procedures were carried out in accord with the UK Animals (Scientific Procedures) Act (1986) and licensed protocols were approved by local committees (LERP and ACER, University of Oxford) and carried out under license number P6CE80C21D. Throughout the procedures, all efforts were made to adhere to the ARRIVE and IMPROVE guidelines (19, 20). Briefly, Animals were randomly assigned to experimental groups prior to EV injection to ensure unbiased treatment allocation. EVs were isolated, labelled, and coded by an independent researcher not involved in the injections or downstream analyses. Group identities were recorded in a central spreadsheet and kept blinded from the experimenter performing the injections and analyses. Blinding was maintained throughout data collection and analysis. For histological assessments, microscope slides were anonymised by covering sample identifiers with opaque tape. Quantification was performed without access to treatment information, which was only unblinded after all data had been collected and analysed.

### Middle Cerebral Artery Occlusion (MCAO) induction

For surgery, animals were anesthetized in 1.5-2% isoflurane carried in an oxygen:nitrous oxide mix (60:40, 2 L/min). Animals were initially placed prone on a nose-cone and a small incision was made over the area of the cortex supplied by the MCA. The skull was slightly thinned and a laser doppler probe was placed to ensure a minimum 70% drop in baseline blood flow occurred at the time of MCA occlusion. At this point, animals were placed supine on the nose cone and a midline incision was made in the neck. The common carotid artery was isolated and temporarily ligated. The MCA was occluded using the Longa method.(21) Briefly, the external carotid was isolated and ligated at the distal end. The internal carotid artery was temporarily clamped and an incision made in the external carotid. A 6-0 nylon filament (Duccol, US) was inserted into the external and held in using a temporary suture around the external. The filament was then manipulated into the internal carotid, after removing the clamp, and proceeded along the middle cerebral artery. Laser doppler blood flow was monitored for a significant drop and at this point the temporary suture was tightened to keep the filament in place during occlusion. For animals undergoing temporary MCAO, the filament remained in place for 25 minutes prior to removal and cauterization of the hole in the external carotid artery and removal of the ligature around the common carotid. For animals undergoing permanent MCAO, the filament remained in place and the animals were recovered. Survival times were hyperacute (2 hours) or chronic (7 days). For those animals surviving for 7 days, gel-based food and additional s.c. saline were provided throughout the recovery period.

### Modelling inflammation

Animals received a single dose of LPS (0.5mg/kg) i.p. to induce a typical systemic inflammatory response (22). Animals were allowed to survive for 6 hours prior to culling when blood was collected for EV isolation.

### Extracellular vesicle (EV) isolation

For all animals, blood was collected by cardiac puncture using a heparinized needle and taken into heparinized tubes. Whole plasma was centrifuged for 15 min at 2500*g*, the supernatant was then centrifuged for a further 15 min at 2500*g*. Platelet-free plasma (PFP) was then aliquoted into 100µl aliquots and stored at -80°C until EV isolation. EVs were isolated fresh prior to each experiment. Briefly, 100µl of PFP was applied to single-use size exclusion columns (70nm Legacy - iZON, UK) and 12 x 200μl fractions were collected, with vesicles being pooled from fractions 1-4 (supplementary Fig.S1). Size exclusion was used to enrich for small extracellular vesicles (EVs), which are likely to predominantly include exosomes and small microvesicles based on size. However, this isolation method may also capture other vesicle subtypes, and does not allow complete discrimination between EV populations – hence the use throughout the manuscript of the umbrella term ‘extracellular vesicle’ (11).

### Extracellular vesicle analysis

Counts were performed using the Zetaview platform (Particle Metrix, UK). EV samples were diluted in PBS (1:1000) and were analysed for size and concentration using NTA video capture for 0.5 s at 11 positions with the following acquisition settings; shutter speed = 100, sensitivity = 80, cycle number = 2, frame rate = 30, minimum brightness = 25, maximum size = 1000, minimum size = 5, trace length = 15 and were analysed using the integral Zetaview software. After counting aliquots of EVs were stored at -80°C until further use.

### Electron microscopy

For negative staining, freshly glow discharged (15 mA, 25 seconds) 300 mesh carbon-coated EM grids (C267, TAAB) were inverted carbon-side down onto a 10 µl droplet of sample and incubated for 2 minutes. Grids were then gently blotted with filter paper and transferred onto a 20 µl droplet of 2% (w/v aq.) uranyl acetate solution. Samples were incubated for ten seconds, blotted and air-dried. Negative staining was kindly performed by Dr Errin Johnson (Dunn School of Pathology, University of Oxford).

### Flow cytometry

The MACSPlex EV kit (IO – immuno-oncology; Miltenyi, UK) was used to not only characterize the surface profile of the EVs, but also to ensure the expression of ‘standard’ markers of EVs (CD63, CD9, CD81), as per MISEV guidelines (12, 23). EVs were isolated from plasma, as above, and 10^12^ EVs per sample were run according to the manufacturers’ instructions with an overnight incubation at 4°C to increase specificity. Samples were run on the CytoFLEX flow cytometer (Beckman Coulter) with lasers adjusted for use with this kit (assistance kindly provided by Dr Josh Welsh and Dr Jamie Cooper). Samples were set to record at least 5,000 single-bead events per sample. Bead populations were identified based on their fluorescence in the PE and FITC channels. For analysis, samples were normalized to background fluorescence units acquired for buffer and isotype controls. The values were normalised to the mean MFI of the expressed markers to determine the relative expression of each marker.

### Dot blots

EVs were isolated from plasma, as above, and 10^12^ EVs were applied to Proteome Profiler XL Array (Mouse; Biotechne, UK) and samples were run according to manufacturers’ instructions with an overnight incubation at 4°C to increase specificity. Image analysis was performed using the circle tool in the BioRad ImageLab software, where a circle of a specific size was drawn and then placed over each spot on the blot (Supplementary Fig. S2). Reference spots for each blot were set as ‘100’ and all dots within that blot were calculated as relative to those, in order to reduce inter-blot variability that might be introduced by comparing blots directly to naïve samples. Relative levels for each blot were then plotted as a heat-map using R.

### Stereotaxic surgery

Animals were anaesthetized (using the same parameters as for MCAO surgery) prior to being placed on a stereotaxic frame. A midline incision was made on the scalp and a burr hole drilled in the skull above the co-ordinates for the dorsal striatum (Bregma +0.5 A/P, -1.5 M/L and -2.5 D/V). 1µl EVs (1×10^10^/µl) in PBS was injected over five minutes using a glass microcapillary. Animals were allowed to survive for 24 hours prior to culling.

### Tissue collection

For fresh tissue, animals were perfused with ice-cold 0.9% saline containing 5U/ml heparin (Sigma, UK). Liver and brain tissue was snap frozen and stored at -80°C until further use. For fixed tissue, animals were perfused with room temperature 0.9% saline until tissues were cleared, followed by 4% paraformaldehyde (PFA) in 0.1M phosphate buffer. Tissue was post-fixed overnight in 4% PFA and then cryoprotected in 30% sucrose until the tissue had sunk.

### Tissue processing

Fresh and fixed brains were cryosectioned at 12μm using a Leica cryostat +/-500μm A/P of the injection location. For fixed sections, tissue was placed on gelatin coated slides, air-dried and stored at -80°C. For fresh section for qPCR, sections were split into ipsilateral and contralateral hemispheres and stored at -80°C.

### Immunohistochemistry and analysis

To assess the naïve CNS response to exogenous EVs, brain sections were stained for intracellular adhesion molecule (ICAM)-1 (1:1000; Sigma-Aldrich/eBioscience 12-0549-42), glial associated fibrillary protein (GFAP; 1:500; AbCam ab7602) ionized calcium binding adaptor molecule 1 (Iba1)+ microglia (Iba-1; 1:500; AbCam; ab178847) with a cresyl violet counterstain. Nonspecific binding was blocked using 10% serum from the species in which the secondary antibody was raised (diluted in PBS) for 1 hour at room temperature (RT) and incubated with the primary antibody in PBS at 4°C overnight. Secondary antibodies were biotinylated (1:100; VectorLabs, UK) and applied for 1 hour at RT followed by visualisation using the avidin-biotin-peroxidase method (1:100, 1 hour at RT, VectorLabs, UK) using 3,3′-diaminobenzidine to visualize the stain. Quantification of immunohistochemical staining was performed using a modified version of a previously published protocol (24). Four representative coronal sections were selected per brain, and the striatum was imaged bilaterally (ipsilateral and contralateral) at 20× magnification using ScanScope software and a Leica Aperio Slide Scanner. Images were processed in ImageJ, where colour deconvolution was applied to separate the DAB signal from cresyl violet (when both stains were present). A square region of interest (ROI) with a fixed diameter of 500 ImageJ units was placed within the striatal region of each image, the location of the quantified region is indicated in the inset panel of Figure 6A. The ‘Measure’ function was used to quantify average staining intensity and percentage area stained (reported in arbitrary units). This analysis was focused on a defined ROI within the striatum rather than the entire hemisphere or brain, and therefore the percentage values reported reflect the area stained within that specific ROI. This technique was applied because ICAM staining is not specific to the vasculature, but can be present on microglia as well and as such traditional cell counting methods are not necessarily appropriate. We therefore used the same technique across all staining moieties.

### qPCR

RNA was extracted using the RNEasy Mini Kit (Qiagen, UK) with QiaShredders. cDNA was generated using the High-Capacity cDNA Kit (Applied Biosystems, UK) and diluted to a final concentration of 5ng/ul. 25ng of cDNA was used in the quantitative PCR reaction with PrecisionPLUS SYBR green master mix (Primer Design Ltd., UK). Primer sequences (5’3-‘) were as follows: GAPDH F: AACGACCCCTTCATTGAC R: TCCACGACATACTCAGCAC; IL-1β F: CAACCAACAAGTGATATTCTCCAT R: GGGTGTGCCGTCTTTCATTA; CXCL-1 F: GCAGACAGTGGCAGGGATT CXCL-1 R: GTGGCTATGACTTCGGTTTGG; ICAM-1 F: CAATTTCTCATGCCGCACAG R: AGCTGGAAGATCGAAAGTCCG; VCAM F: TGAACCCAAACAGAGGCAGAGT R: GGTATCCCATCACTTGAGCAGG.

### Statistical analysis

Statistical analyses were performed using GraphPad Prism 9.0 software. All data were tested for normality and post-hoc tests adjusted accordingly. Results were considered statistically significant at p<0.05.

## Results

### Neither stroke nor systemic inflammation induce significant increases in circulating EV numbers

EVs were isolated from the supernatant of either animals after middle cerebral artery occlusion (MCAo) where the filament was removed after 20 minutes (transient occlusion – tMCAo) or where the filament remained in place (permanent occlusion – pMCAo). Animals were allowed to recover for 2 hours (acute) or 7 days before plasma was collected for EV isolation. These animals were compared to those who had experienced an acute inflammatory challenge (6h i.p. LPS), to determine the degree to which the circulating EV population might reflect an inflammatory response. After size exclusion chromatography (SEC), the first four fractions, where contaminating free proteins were excluded (Supplementary Fig. S1), were pooled and analysed by nanoparticle tracking. There was no significant difference in average particle concentration between the groups (Fig.1A). Distribution suggests that EVs lie within the normal range expected (Fig. 1B-C) and that reperfusion of the brain does not appear to significantly influence the distribution pattern of EVs found in the plasma. Electron microscopy shows the standard ‘cup-shaped’ EVs lying between 50-200nm (Fig. 1B-C - insets), demonstrating that SEC of cell culture supernatant is an effective way to isolate EVs. However, electron microscopy images also show some ‘EV-like’ structures and an abundance of miscellaneous particles (Fig.1B-C – inset). These are likely lipoprotein contaminants and their removal was beyond the scope of the current study.

**Figure 1.**
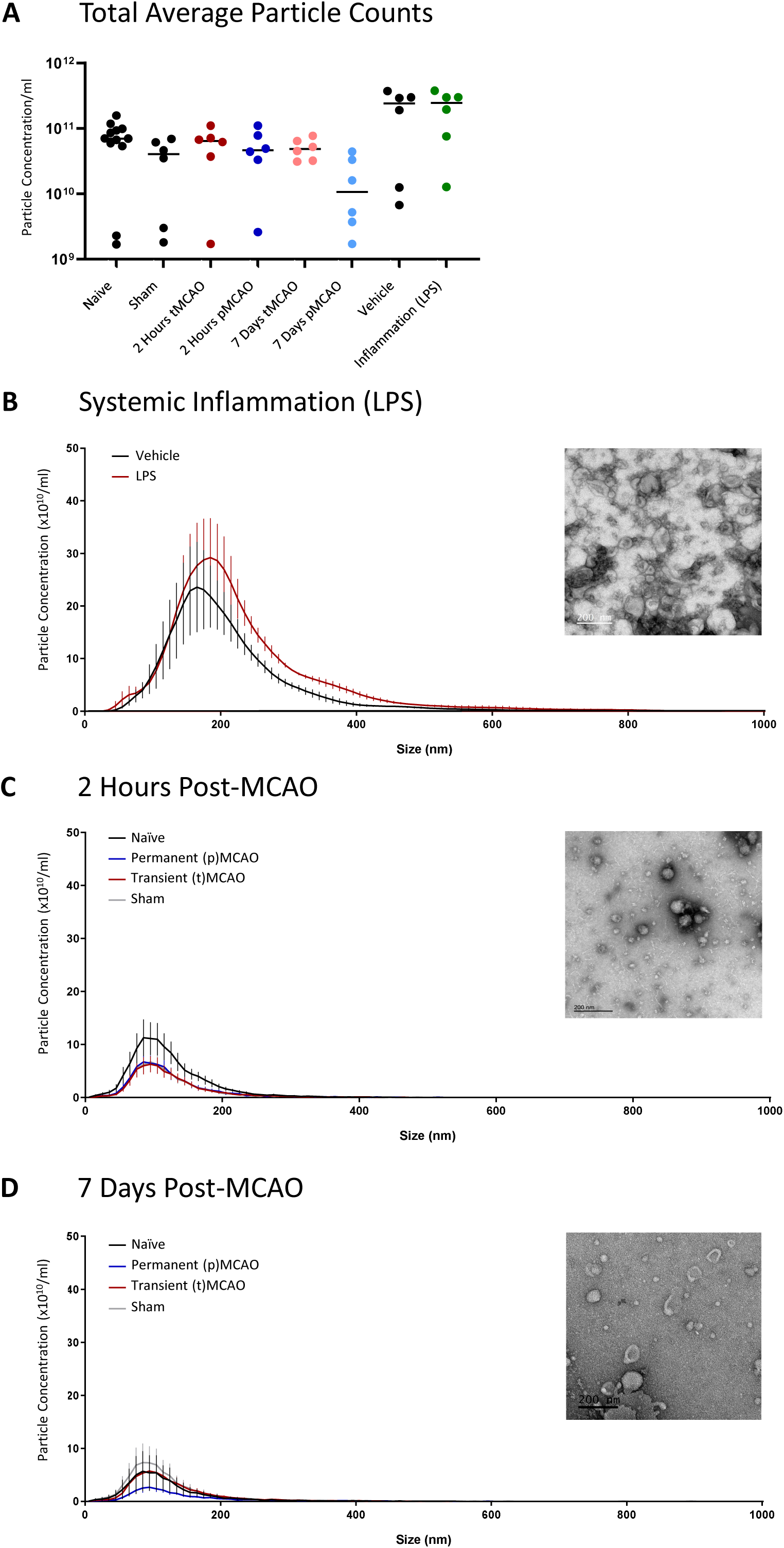
Extracellular vesicle characterization by dynamic light scattering and electron microscopy in models of acute stroke with and without reperfusion, and systemic inflammation. Animals were subjected to either transient middle cerebral artery occlusion (tMCAO) where the filament was removed from the middle cerebral artery after 25 minutes, or permanent middle cerebral artery occlusion (pMCAO) where the filament remained in place. Sham animals underwent similar surgery but the filament was not advanced into the middle cerebral artery in any cases. All stroke animals were allowed to survive for 2 hours or 7 days. Inflammation was modelled using an intraperitoneal injection of lipopolysaccharide (LPS) and animals were allowed to survive for 6 hours. Plasma was harvested and EVs isolated from platelet-free plasma using size exclusion chromatography (SEC) followed by analysis using the Zetaview platform and by negative staining electron microscopy. (A) Total average particle counts do not change significantly between groups. (B) Size distribution and electron microscopy of EVs from animals post-LPS challenge. EVs are within the normal range for EVs isolated by SEC and electron microscopy shows some standard ‘cup-shaped’ EVs. Similar data for animals at 2 hours post-MCAO (C) and 7 days post-MCAO (D) with and without reperfusion. EVs show similar characteristics. Data are individual animals with the mean indicated, n=6-12.

### Mouse plasma EVs express high levels of integrins and adhesion molecules, as well as CD9 and CD81 but low levels of CD63, irrespective of treatment

Given the small starting sample volumes for mouse plasma experiments, the use of traditional techniques such as Western blotting (12, 23) for confirming the presence of EVs was challenging due to low protein yield. As such, more specialist techniques such as dedicated flow cytometry panels, were used. Here, the EV-specific mouse immuno-oncology panel includes the tetraspanins CD9, CD81 and CD63 and should be considered a surrogate for blotting. All samples, including naïve, showed high expression levels of CD9 and CD81 suggesting that EV isolation was successful and that the fractions chosen from early SEC did indeed contain EVs (Fig.2, arrows). No samples showed significant expression levels of CD63 (Fig.2, arrows). In addition to standard markers of EVs, early SEC fractions also showed increased expression levels of integrins such as CD62P, CD29 and CD61 as well as inflammation-associated molecules such as MHCII and CD44. As expected, given the high prevalence of platelet EVs in plasma samples, all samples also expressed high levels of CD41 (Fig.2). Although individual one-way ANOVAs identified some nominal differences in specific markers, these were not emphasized due to the risk of inflated Type I error and absence of correction for multiple comparisons (raw data available in *Supplementary File SF1*).

**Figure 2.**
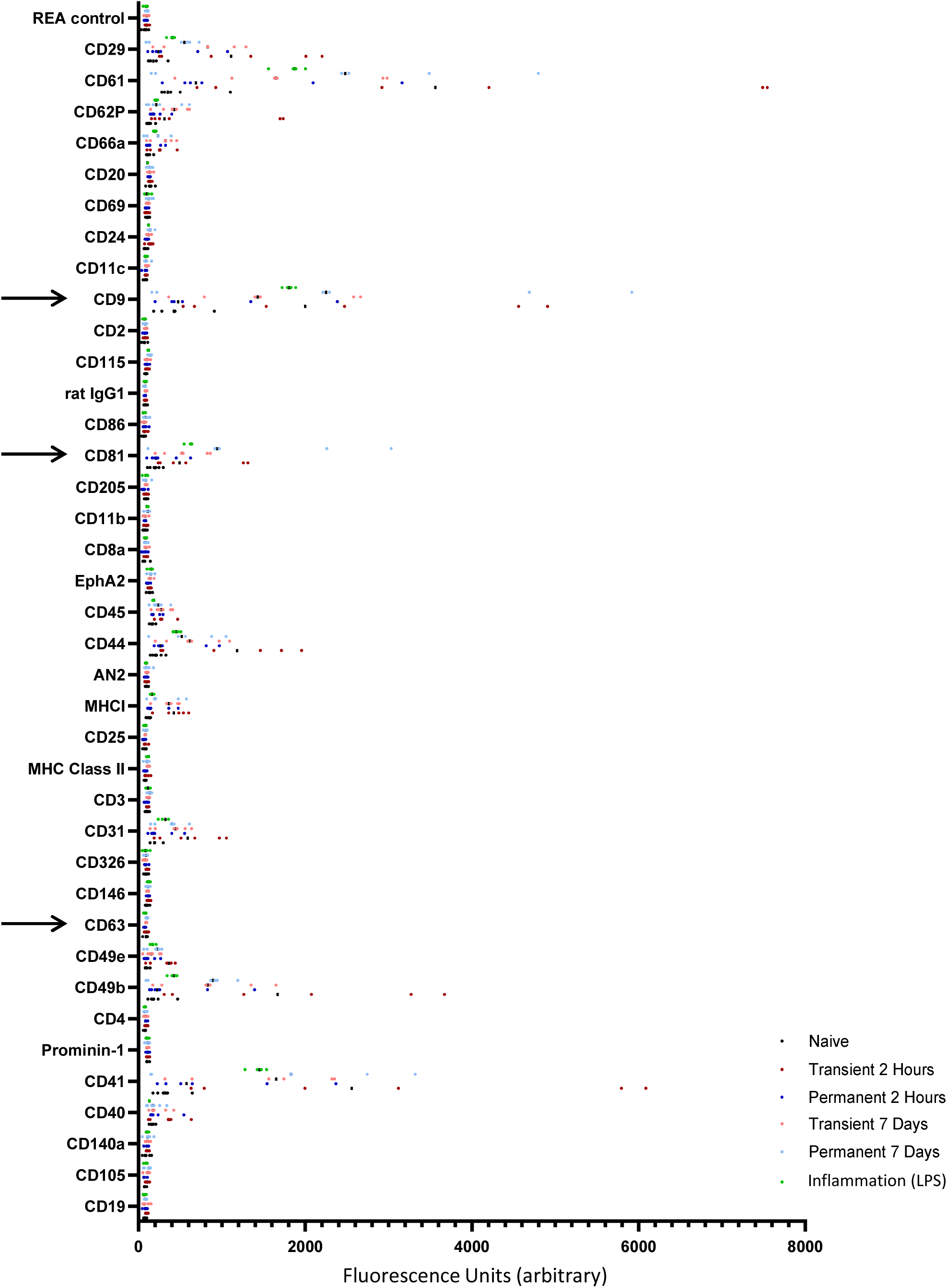
Flow cytometric characterization of extracellular vesicles in models of ischemia and inflammation using the immuno-oncology EV panel. After isolation using SEC, EVs were incubated with a panel of bead-bound membrane marker antibodies in combination with a cocktail of bead-bound tetraspanin antibodies (CD9, CD63, CD81) and run on a modified flow cytometer. EVs show characteristically high levels of tetraspanins, demonstrating successful EV isolation, highlighted here with arrows at CD81, CD9 and CD63. Data are individual animals with the mean indicated, n=4.

### Reperfusion of the brain results in EVs in the circulation with a more pro-inflammatory protein profile

The preservation and amplification of signal on EVs is a known mechanism of organ-to-organ communication.(25, 26) Here, we wished to investigate the potential for the protein profile of EVs in the circulation to be affected by the original pathological stimulus. Given that inflammation is a known post-stroke outcome we used a prototypical inflammatory stimulus (LPS) to generate plasma EVs with a stereotypical inflammatory profile. The data show that pooled EVs from the plasma of animals undergoing inflammation show upregulation of inflammatory markers on their EVs (Fig.3). These include C-reactive protein (Fig.3, highlighted box 1), E-selectin (Fig.3, highlighted box 2) and pentraxin-3 (Fig.3, highlighted box 3). These proteins are also present at high levels in EVs from animals two hours post-MCAO when the brain has been reperfused, but appear lower in those animals where the brain has not been reperfused. E-selectin, for example, is particularly high at two hours post-tMCAO but remains low post-pMCAO until the seven-day timepoint, where it is possible chronic inflammation may be starting. Other molecules, such as serpin-E1, (Fig.3, highlighted box 4), appear only in prototypical inflammation but are not as highly expressed in any of the post-stroke EV samples (raw data available in *Supplementary File SF2*).

**Figure 3.**
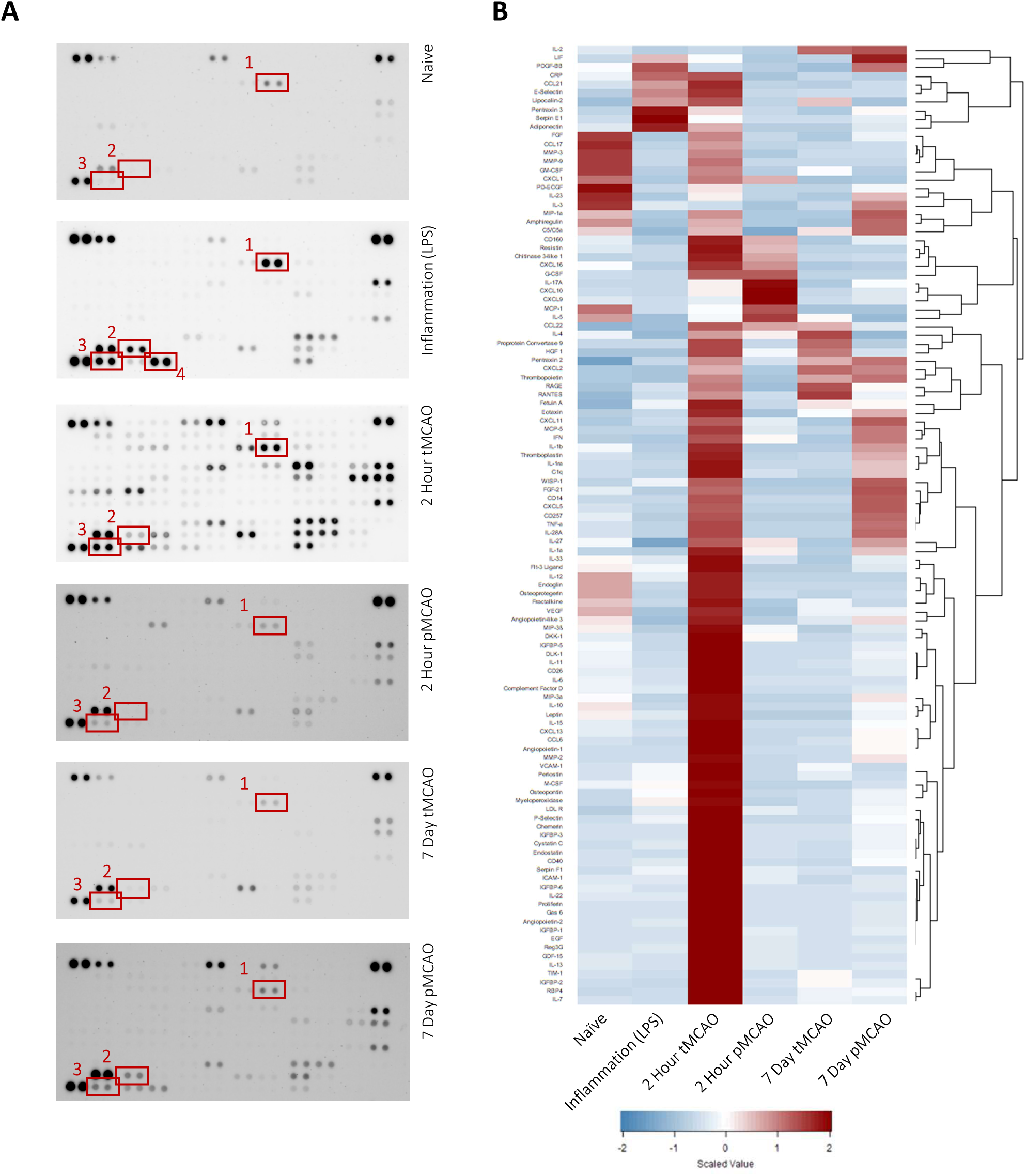
*Differential cytokine and chemokine expression in extracellular vesicles in models of ischemia and inflammation*. After isolation using SEC, EVs from 4 animals were pooled and were applied to the pre-labelled Proteome Profiler membrane and developed according to the manufacturers’ instructions. Dot blots from naïve, inflammation, 2-hour tMCAO, 2-hour pMCAO, 7-day tMCAO and 7-day pMCAO-derived EVs are shown (A). Expression levels within blots were normalized relative to the pairs of control dots (top left and right corners, bottom left corner) and then relative expression levels were compared across groups (B). Boxes show changes in specific cytokines and chemokines across groups which may be particularly notable. These include (1) C-reactive protein, (2) e-selectin, (3) pentraxin and (4) serpin which is highlighted as being particularly high in the inflammation-derived EVs only.

### EVs with different pathological origins induce different inflammatory responses

Given the differences in protein profile on the EVs from different sources, our assumption was that they would have different physiological effects. One way which EVs might be important post-stroke is through the communication of inflammatory signals to the circulation, but also to distant parts of the brain. Here, we used a glass microcapillary to inject EVs into the naïve brain to determine their effect on the local immune cell population. In particular, we were interested in the degree to which pro-inflammatory genes were locally upregulated in response to exogenous EVs. We found that EVs derived from animals post-stroke with and without reperfusion of the brain induced different inflammatory profiles depending on when the EVs were harvested post-injury. Interestingly, despite the increased pro-inflammatory profile from transient EVs, they did not induce significant expression of either IL-1 or CXCL1 after injection into the naïve brain (Fig.4A-D). EVs derived from animals with a permanent occlusion of the MCA did induce significant upregulation of IL-1 and CXCL1 at 2 hours post-occlusion (Fig.4A &B; p<0.05, p<0.001 respectively) but only of CXCL1 at 7 days post-occlusion (Fig.4D; p<0.0001). We also found that EVs derived from the circulation of inflammatory animals induced inflammatory gene expression in the brain. However, they – like the EVs from transient occlusion animals, did not upregulate IL-1 (Fig.4E) but did upregulate CXCL1 mRNA at the site of injection (Fig.4F; p<0.01).

**Figure 4.**
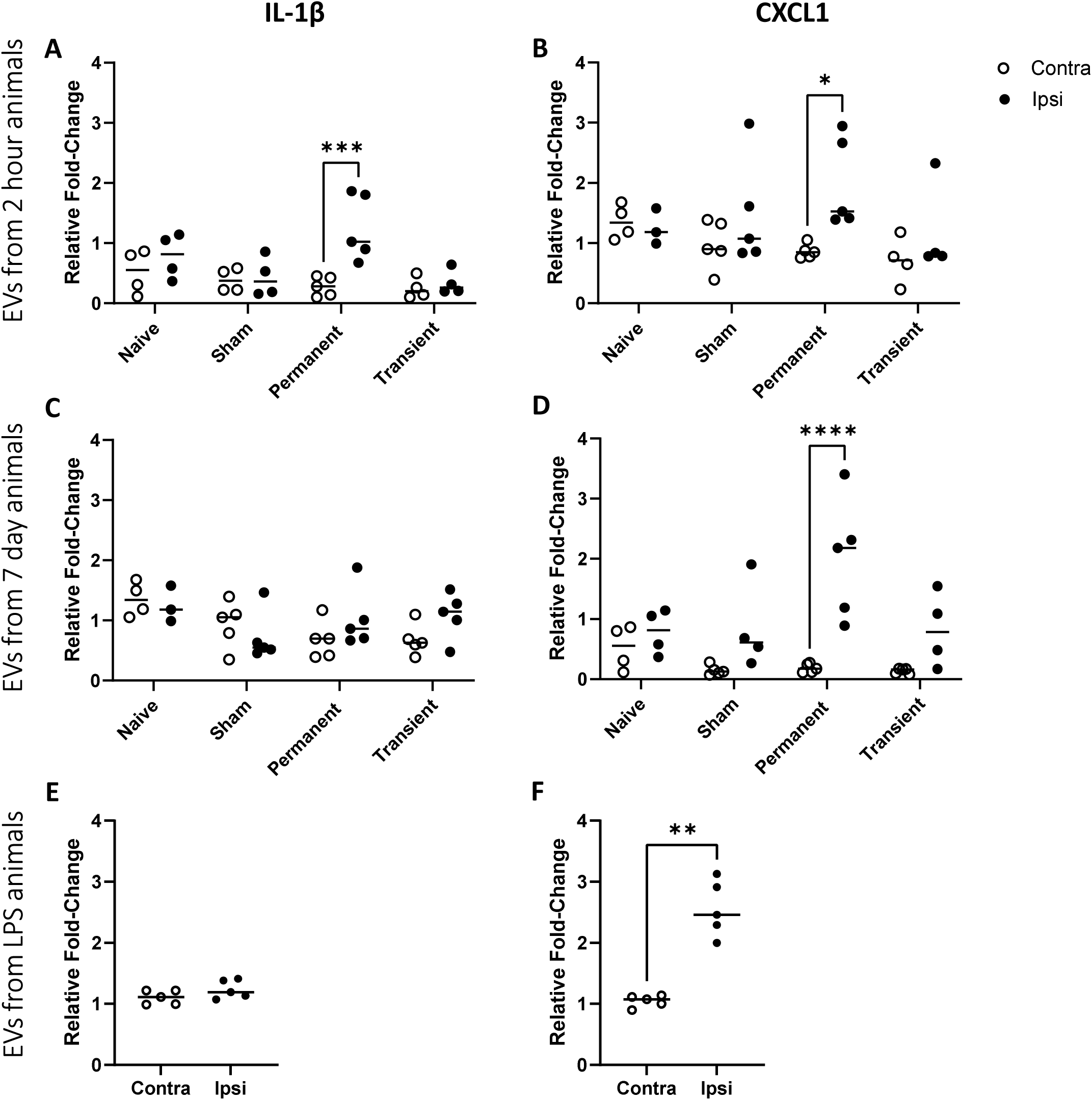
Proinflammatory cytokine and chemokine mRNA expression in the brain after injection of EVs derived from models of stroke and inflammation. After isolation using SEC, EVs from all groups were quantified and 10^10^ EVs from each group were injected in 1µl into the striatum. Tissue was snap frozen isolated striata were processed for mRNA. qPCR of pro-inflammatory cytokine IL-1β and pro-inflammatory chemokine CXCL-1 demonstrate some increases in the ipsilateral hemisphere after injection of EVs. EVs derived from different initial challenges resulted in different expression profiles. EVs from animals 2 hours post-stroke increased IL-1 (A) and CXCL-1 (B) expression only after permanent occlusion. EVs from animals 7 days post-stroke did not increase IL-1 expression in any model (C) and increased CXCL-1 only after permanent occlusion (D). EVs from animals after an LPS challenge did not increase IL-1 (E) but did increase CXCL-1 expression (F). Data are individual animals with mean indicated, n=4-5. *p<0.05, **p<0.01, ****p<0.0001.

Given the presence of markers of inflammation, such as CRP, on the surface of our EVs (Fig.3) and the fact that these inflammatory molecules are known to upregulate endothelial adhesion molecules (27), we hypothesized that our EVs would likely upregulate ICAM and VCAM in naïve animals. In fact, irrespective of which animals or time-points our EVs were derived, they induced no significant upregulation of VCAM expression in the brain (Fig.5A, C & E). However, EVs from animals experiencing a transient MCA occlusion upregulated ICAM mRNA expression in the brain when taken at both two hours and seven days post-occlusion (Fig.5B & D; p<0.05 and p<0.01, respectively). Similarly, EVs derived from animals after acute inflammation (LPS) also upregulated ICAM mRNA expression when injected into a naïve brain (Fig.5F; p<0.05).

**Figure 5.**
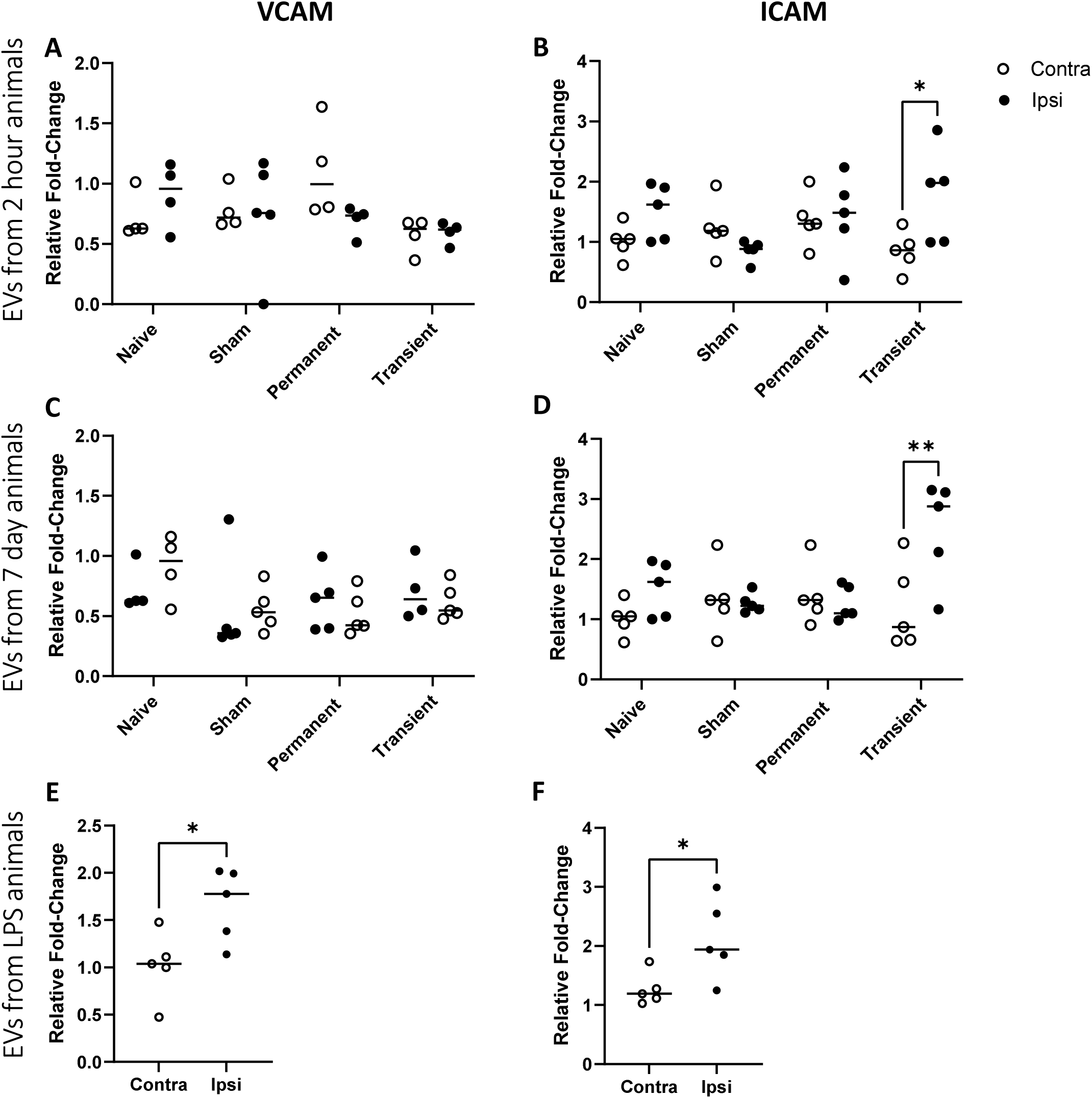
*Adhesion molecule mRNA expression in the brain after injection of EVs derived from models of stroke and inflammation*. After isolation using SEC, EVs from all groups were quantified and 10^10^ EVs from each group were injected in 1µl into the striatum. Tissue was snap frozen isolated striata were processed for mRNA. qPCR of vascular adhesion molecule-1 (VCAM) and intracellular adhesion molecule-1 (ICAM) demonstrate some increases in the ipsilateral hemisphere after injection of EVs. EVs derived from different initial challenges resulted in different expression profiles. EVs from animals 2 hours post-stroke did not increase VCAM expression (A) but did increase ICAM expression (B), the latter only after transient occlusion. Similarly, EVs from animals 7 days post-did not increase VCAM expression (C) but did increase ICAM expression (D), again only after transient occlusion. EVs from animals after an LPS challenge increased both VCAM (E) and ICAM (F) expression. Data are individual animals with mean indicated, n=4-5. *p<0.05, **p<0.01.

### EVs with different pathological origins induce activation of different cell types in the brain

In addition to mRNA expression profiles indicating localized inflammation, it was important to determine whether there was histopathological evidence of localized inflammation. To this end, tissue was stained for markers of microglial activation, astrocyte activation and vascular inflammation and tissue was analysed in the ipsilateral hemisphere only (see Supplementary Fig.4 for 20x image of cells and for images of whole brains to demonstrate the hemispheric differences where present). ICAM expression was increased in animals injected with EVs from 2-hour transient (Fig.6A & D; p<0.01) and 2-hour permanent (p<0.05) stroke animals, similarly in those animals receiving EVs from the model of inflammation (Fig.6A&D; p<0.01). GFAP expression did not change across treatment groups when the area of injection (the striatum) was analysed (Fig.6B & E). However, there was a significant difference in adjacent GFAP reactivity in the corpus callosum (supplementary Fig.3A & B). Similarly, Iba-1 expression looked promisingly prevalent in brains injected with inflammation-derived EVs but this was not significantly different from brains injected with EVs from any other origin (Fig.6C & F).

**Figure 6.**
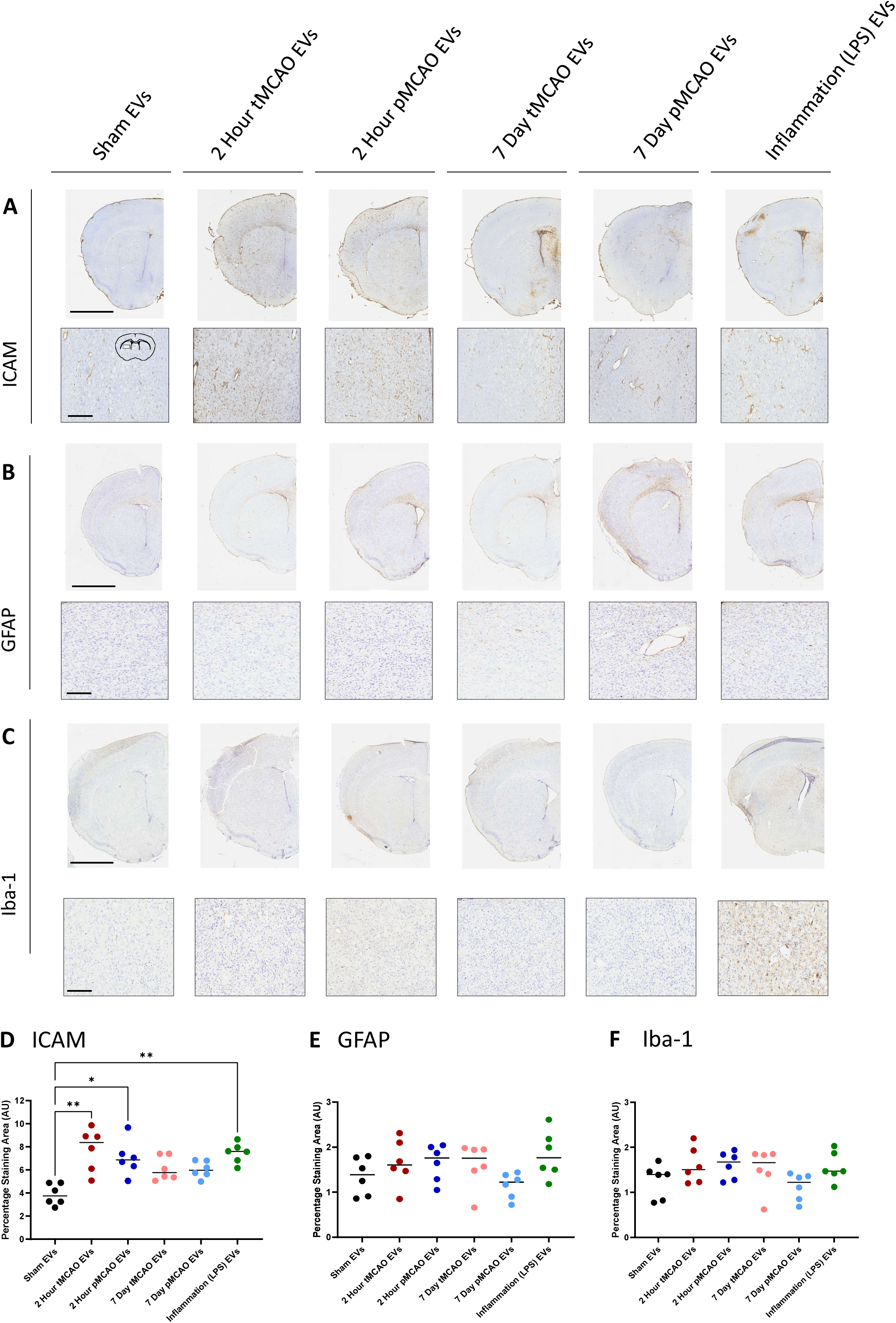
ICAM, GFAP and Iba-1 expression in the brain after injection of EVs derived from models of stroke and inflammation. After isolation using SEC, EVs from all groups were quantified and 10^10^ EVs from each group were injected in 1µl into the striatum. Tissue was fixed and cryosectioned and stained for ICAM (A), GFAP (B) and Iba-1 (C). All tissue was also counterstained with cresyl violet. Example images are of the ipsilateral hemisphere and a magnified area showing detailed staining in each case, coronal section outline in sham panel in (A) shows the area selected for the higher magnification image. Animals injected with EVs from sham animals show very little expression of either ICAM or GFAP, EVs derived from models of stroke and inflammation show varying levels of expression of ICAM (D) but less variability in expression levels of GFAP (E) or Iba-1 (F) when the striatal staining area was analysed. Scale bars on images of hemispheres represent 1mm and scale bars on images of higher magnification represent 200um. Data are individual animals with mean indicated, n=5. *p<0.05, **p<0.01.

## Discussion

EVs are known to be a means of cell-to-cell communication but their role in the potentiation of injuries such as stroke is currently unclear. Here we have demonstrated that the original injury contributes to the physical make-up of the EV, and that these different characteristics result in different inflammatory responses within the brain. Given the increasing interest in using EVs as biomarkers in diseases such as stroke (28), this paper demonstrates the need to approach these studies with caution, making sure that a diverse patient population with differing pathological ӕtiologies is represented, in order to extrapolate effective biomarkers of disease progression (17). It also demonstrates the paucity of data currently available on the characterization of circulating EVs in mice. In our pursuit of truly bench-to-bedside research it is important, going forward, to consider the differences between commonly used laboratory species and ourselves (9, 10).

At the start of the field of EV research it was assumed that stimulation of any kind, be it in cell culture or as part of a disease state, resulted in increased EV production in the supernatant or bodily fluid they were being measured in (29). However, it is now more widely accepted that absolute particle measurement from biofluids is challenging given the degree of lipoprotein contamination and the challenges associated with isolating a ‘pure’ population of EVs (17, 18, 23). Indeed, studies in stroke show widely varying correlations between infarct volumes, outcomes and EV numbers with studies showing a positive correlation (30) and no correlation at all (31, 32). These differences are likely to reflect differences in isolation and analysis techniques that are ubiquitous throughout the EV field and are the main reason behind the lack of correlational analysis between lesion volume and EV numbers in this study.

Here we have demonstrated that stroke does not induce a particularly significant change in EV numbers when compared to either sham animals or naïve animals, and that this is similarly the case with systemic inflammation. This appears in contrast to our own previous findings in acute stroke patient data (13), but this is likely to be an artifact of isolation and measurement techniques (18, 23). Here we have used SEC as a method of isolating EVs, something which is known to potentially reduce the overall number of ‘particles’ measured by techniques such as dynamic light scattering, but which improves the particle:protein ratio and thereby the purity of the EV sample (33). This can manifest as higher EV counts per mL compared to ultracentrifugation methods, especially when combined with optimized acquisition settings. Additional factors—such as species differences (mouse vs. human), sample handling, and NTA parameters—further complicate direct comparisons with prior work. Furthermore, Pieter Vader’s lab have demonstrated that EVs isolated by SEC seem to have more functional capacity (34), potentially because EVs isolated by UC are more likely to aggregate and rupture. In this study, there seems to be a spread of size distribution with the inflammatory EVs appearing slightly larger than the stroke EVs. However, the naïve group from the LPS animals also show a larger size than the stroke groups. Others have demonstrated significant differences in size patterns *in vitro* (35, 36) and effects of storage on size distribution (37), and as such this difference is considered within the scope of normal experimental and biological variability.

The EVs isolated here express high levels of CD9 and CD81 but not CD63. This has been found elsewhere in cell lines (38) and studies on plasma and serum have found that plasma often has significantly lower levels of CD63+ EVs than serum (39). Given that there may also be a number of differences in EV marker levels between species (40, 41) it is possible that CD63 might not be the ideal marker to identify EV populations in mouse plasma. Indeed, other studies using a similar kit report higher levels of CD9 and CD81 suggesting there may be differences in the basal expression levels of tetraspanins between mice and humans and as such the poor expression levels of CD63 found in this study should not be considered exclusionary for successful EV isolation (42).

In addition to standard surface markers, EVs from all animals expressed high levels of CD41, suggesting a strong platelet origin.(43) The data also demonstrate the inherent noise in the stroke models used, with significant variability present in all post-stroke animals but absent in naïve and LPS-treated animals. The tight clustering of EVs from a model of inflammation does not always show the same pattern of surface marker expression as EVs from post-stroke animals, suggesting that post-stroke inflammation and TLR-driven inflammation do not generate the same EV populations. For example, CD41 expression is variable and high in the acute transient stroke animals, but much tighter - although still elevated – in the LPS animals. Given the important differences in immune-thrombosis and thrombo-inflammation (44), these differences may be key to understanding the origins and purpose of the inflammatory response post-stroke.

This difference is further backed up by the absence of some markers of inflammation on EVs derived from post-stroke animals. PAI-1 (serpin-E1), for example, is induced by circulating cytokines and is expected to be significantly upregulated after LPS stimulation (45). Indeed, the work here shows it to be increased on inflammation-derived EVs but in relatively low abundance in most post-stroke EVs. In contrast, C-reactive protein (CRP), which we have previously shown to be increased on stroke-derived EVs (13), is present on both inflammatory EVs and on stroke-derived EVs, with higher abundance earlier in the disease process. This is reflected by standard measures of plasma CRP levels after brain injury, which show peaks in the early phases and a decline over time (46). Whilst this may suggest that EV-derived CRP might be used as a prognostic biomarker, it could be argued that CRP levels can be easily measured in the plasma without the additional technical challenges of EV isolation (17) and as such this may not add a great deal to our capacity for patient stratification. However, we do know that EV surface characteristics change the physical properties of the CRP pentamer (25), meaning that whilst it may not be a useful biomarker, understanding the role of EV-bound CRP in post-stroke inflammation and recovery continues to be an important question which was, sadly, beyond the scope of the current study.

There is much that continues to be poorly understood about EV biology, especially *in vivo*. One pressing question is the purpose for the EV-expression of specific proteins. Whilst the stochiometric changes of CRP could be seen to serve a specific, pro-inflammatory purpose, the release of molecules such as E-selectin on EVs remains more complex. Early studies, around the time EVs were beginning to break into the literature, found that E-selectin was released from activated endothelial cells (47). The authors suggest that release of soluble adhesion molecules, irrespective of mechanism, could be a means of inducing local immune cell activation, or a means of inhibiting immune cells’ adhesion to the vasculature by locally blocking the receptors.

The inflammatory response to ischemia with reperfusion vs without reperfusion has been documented elsewhere (48, 49). Indeed, changes in inflammatory activity and vascular reactivity between the two models are likely to be responsible for the differences in EV populations observed here. Previous work has shown that reperfusion of the CNS results in significantly different expression levels of vascular adhesion molecules (50). Furthermore, studies of shear stress, a phenomenon known to be different between the two models, demonstrate that changes in blood flow affect EV release from endothelial cells (51). In addition to the physiological differences between the models, some degree of difference in EV populations is also expected between time-points. This study included acute and chronic time points based on established time windows in stroke pathophysiology, where key phases of the inflammatory response are known to occur. At 2 hours post-stroke, the acute phase of inflammation is initiated, and this timepoint captures the early events of blood-brain barrier disruption, microglial activation, and infiltration of peripheral immune cells. At 7 days post-stroke, there is typically a shift toward chronic inflammation, with ongoing activation of astrocytes and microglia, and the onset of tissue repair processes with no obviously visible absence of cresyl type-lesion (52). This timepoint was chosen to reflect a later, more stable phase of the inflammatory response, allowing us to assess the potential persistence of EV-associated inflammatory changes. Detailed flow cytometric analysis of adhesion molecules on EVs would provide more insight into their potential origins and the temporal effects of stroke and reperfusion, but were, unfortunately, beyond the scope of this study.

Beyond the use of EVs as diagnostic or prognostic biomarkers, it is important to understand their role in physiology and pathology (9, 10). Here, introduction of EVs into the CNS had a variety of effects. Significant activation of the vasculature was found (ICAM expression), as well as increased GFAP expression within the corpus callosum, mostly in response to EVs derived from plasma taken during the acute stages of reperfusion, or from inflammation. The paucity of VCAM and Iba-1 staining in response to these challenges was puzzling and remains the focus of ongoing studies. However, some initial speculations can be made.

The roles of ICAM and VCAM in the brain are separated by sensitivity and expression. ICAM is much more ubiquitous, being expressed on astrocytes and microglia, as well as endothelial cells, and is one of the first adhesion molecules to be activated during ischemia (53). ICAM is an early warning system and much more sensitive to low levels of inflammation and changes in shear stress (54, 55). It is not unsurprising that ICAM protein expression followed a similar pattern to ICAM mRNA and the response may reflect the low-level inflammatory potential of EVs from LPS-challenged animals, and from those deriving from the plasma of acute stroke animals. It is possible that higher concentrations of EVs or more prolonged exposure may result in increases in VCAM expression and resultant system immune cell migration. These questions remain the focus of our ongoing research.

The absence of microglial reactivity is more perplexing but may be explained by the challenge itself. Astrocytes express lipid receptors which many microglial cells do not. They are sensitive, for example, to levels of LDL and lipoproteins are known contaminants of EV preps from plasma, even when using SEC (12). It is understood that cholesterol may play a role in astrocyte-mediated neuroinflammation (56) which may not occur in microglia. Together with the data on the lack of VCAM signal, this suggests that our EVs may simply not be inducing a significant inflammatory response. Whilst an obvious next step would be to increase the dose of EVs, this might not represent what occurs in normal pathology, where EVs are likely to be released over long periods of time. As such, an osmotic mini-pump approach which continuously infuses low levels of EVs into the CNS might be more appropriate and is the current focus of our ongoing research in this area.

### Study Limitations

The limitations of this are largely the limitations of the EV field generally. Although our isolation approach enriches for small EVs, it does not fully exclude the presence of other extracellular vesicle subtypes such as apoptotic bodies or migrasomes. Future work using orthogonal isolation and characterization methods will be required to comprehensively define the complete EV landscape following stroke, along with more effective tools to distinguish these populations from additional ‘particle’ populations within biofluids. Because whilst size exclusion does remove the majority of protein contaminants, ideally we would also use cation exchange to remove much of the lipoprotein contamination (57) which continues to interfere with ‘particle’ measurements, but this is challenging with the very small volumes of plasma obtained from mouse samples. One possible solution would be to do larger numbers of initial injury models and pool the plasma samples prior to isolation but this would be somewhat against the principles of the NC3Rs (58).

Additional characterization challenges relate to the relative scarcity of *in vivo* data using mouse biofluids, and the consequential lack of good positive and negative markers. This is highlighted by our low levels of CD63, which are not seen in human populations of plasma EVs, where CD63 levels tend to be high. This study also used a multiplex bead-based approach to profile a broad panel of EV surface markers. Whilst this allows for high-throughput comparison across groups, the resolution for individual markers is limited, and subtle or low-abundance differences may be missed. Furthermore, the need to correct for multiple comparisons in high-dimensional datasets reduces the interpretability of isolated marker changes unless independently validated, which would be an important next step in this research.

It should also be noted that, using the techniques in this paper, the majority of our EV population expressed markers that suggest they may be of non-CNS origin. We know that a significant acute phase response occurs post-stroke (59), and that there is immune system activation and changes in platelet behaviour (60), meaning these populations of cells are likely to contribute to the circulating EV population. However, the isolation approach used here does not distinguish between EVs derived from different cellular origins, and thus the signatures observed likely reflect a composite of EVs from multiple tissues. Whilst we did include an ‘inflammatory’ EV group, the acute phase response induced by pathogen associated molecular patterns such as LPS, and that induced by damage associated molecular patterns such as those occurring during injuries, are likely to be different. To expand on this further proteomic mapping of EVs and likely cells of origin would need to be undertaken and a bioinformatics study of this scale was beyond the scope of the current study.

Indeed, whilst this study demonstrates that EVs derived from different sources can elicit divergent effects within the brain, the underlying mechanisms driving these differences remain unresolved. Specifically, we did not attempt to isolate or identify the active cargo components (e.g., proteins, RNAs, or lipids) responsible for the observed outcomes. This reflects both a limitation of the current work and a broader challenge within the EV field. The functional complexity of EVs, coupled with technical constraints around isolating pure subpopulations and attributing bioactivity to specific molecular constituents, makes mechanistic dissection extremely difficult, especially if one were to take into account the fact that EVs are likely to mediate their effects via a combination of surface protein interactions and intravesicular cargo (9, 10, 18, 61, 62). As such, our findings should be interpreted as evidence of source-dependent EV effects, rather than conclusive insights into their biochemical underpinnings. This work has demonstrated that the nature of the initial injury can alter the EV population to the degree that it mediates slightly different functional effects on tissues, but makes no claims as to the biochemical origins of those differences. Future work with targeted molecular profiling and functional assays will be essential to define the precise mechanisms involved.

Another obvious limitation is the lack of evidence of specific cellular uptake within the CNS. It is important to note that in this study, we did not directly assess the localization or diffusion of injected EVs within the brain parenchyma. As such, we cannot determine whether the observed inflammatory changes are due to EV entry into brain tissue, adhesion to the vascular endothelium, or activation of perivascular cell types such as endothelial cells, pericytes, or astrocytes. Current techniques used to label EVs often take advantage of their lipid membrane but there are also obvious pitfalls to this approach, not the least of which being the aforementioned significant lipoprotein contamination (63). We have previously used transduction of EVs with non-mammalian RNA (14) to determine the uptake of EVs in different organs and tissues after systemic injection. Whilst it remains unknown to what degree transduction might affect the protein/lipid structure of EVs (and as such their function), combining this technique with single cell isolations from the CNS to determine whether there is preferential cellular uptake of EVs, form part of our ongoing studies.

It is also important to note that the findings presented here reflect a deliberate focus on the local effects of EVs within the CNS, rather than their potential systemic actions. While the broader immunomodulatory roles of CNS-derived EVs remain an important area of investigation, the current study was designed to assess direct, local responses following intracerebral delivery. Whilst this approach, although it is consistent with previous work in the field studying the effects of neuroinflammatory mediators (14, 24, 64), involves a non-physiological route of administration and does not address *in vivo* kinetics or peripheral immune activation. As such, the conclusions drawn here are specific to CNS-localized responses, and caution should be exercised in extrapolating to systemic or translational contexts. Further work is needed to explore how EVs may behave following more physiological modes of release and circulation, and whether they contribute to immune signalling at remote sites.

## Conclusion

The functionality of EVs, especially *in vivo*, is an under-researched area (9, 10). This is, in part, due to the technical challenges associated with isolating and measuring EVs and, in part, due to the nature of EV release under physiological conditions. In a stroke, for example, EVs are likely to be released into the circulation during the acute phase, but during the recovery phase the pathophysiological processes in the CNS are going to be different, resulting in a different EV population. This means that *the in vivo* responses we get to isolated populations are, given current technological limitations, only ever going to present us with a snapshot of the role of EVs in ongoing pathological processes. In future experiments, using devices that administer EVs more slowly (such as osmotic mini pumps) is likely to reflect the gradual release and uptake of EVs more accurately but may still fail to reflect the complex temporal dynamics of EVs *in vivo*. Despite these technical limitations, the data in this paper demonstrates clearly that the origins of EVs in pathology affects not only their physicochemical composition, but also the downstream effects they have on other cells. Establishing the mechanisms that underlie these processes is vital to our long-term understanding of how pathological events, such as a stroke, can affect the brain.

## Acknowledgements

The author would like to acknowledge all the funding and support provided by first Alzheimer’s Research UK, then by the Oxford British Heart Foundation Centre of Research Excellent. Dr Errin Johnson of the Dunn School at the University of Oxford for her expert running of the electron microscopy of the extracellular vesicle samples. In addition, special thanks go out to Dr Lizzie Dellar for much results chat and tea, to Dr Ryan Pink and Jamie Cooper for many breakfasts and assistance with the Macsplex and to Dr Becky Carlyle for an introduction to the joys of R. Final thanks go to all the EV researchers I reach out to on a semi-regular basis who are willing to read my emails and offer advice and thoughts.

## Author Contribution

The author was solely responsible for all aspects of the study, including conceptualisation, methodology, investigation, data curation, formal analysis, visualisation, and writing—both the original draft and subsequent revisions. The author also performed project administration, secured resources, and approved the final manuscript.

## Data Availability

The datasets generated and analysed during the current study are partially available in the Supplementary Information files. Raw data from dot blot and MACSPlex EV assays are provided as Supplementary Tables. Data supporting the findings of the qPCR and histological analyses are presented in full within the main figures of the manuscript. Additional information is available from the corresponding author upon reasonable request.

## Ethics, Consent to Participate & Consent to Publish

All animal procedures were carried out in accordance with the UK Animals (Scientific Procedures) Act 1986 and EU Directive 2010/63/EU, and were approved by the UK Home Office, and ACER and AWERB ethics committees at the University of Oxford under project license PP7444704 (Couch: Molecular Mechanisms of CNS Injury). Consent to participate and consent to publish are not applicable.

---

**Supplementary Figure 1.**
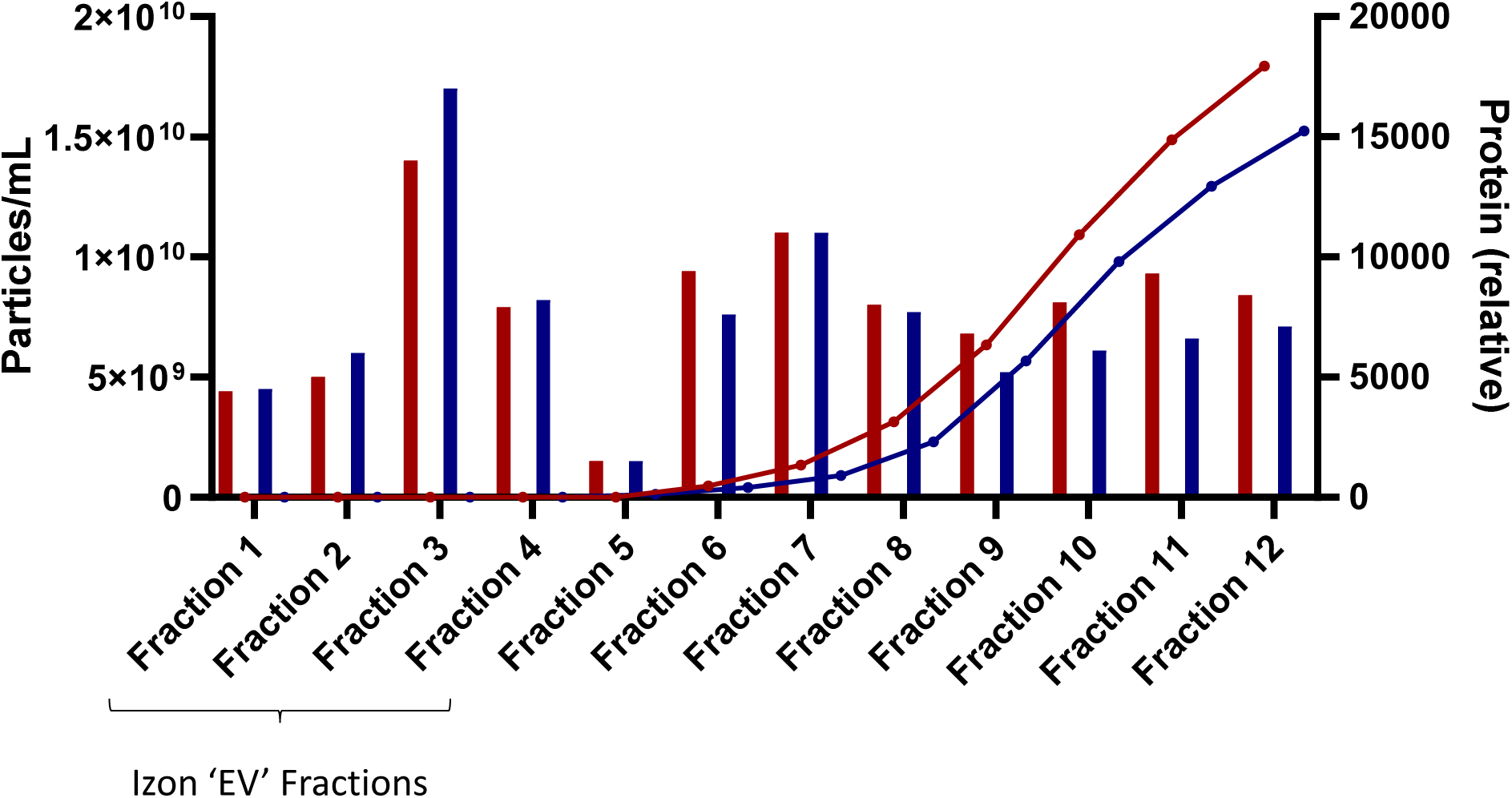
Particle vs protein analysis. In order to determine the degree to which SEC was removing contaminating protein we analysed the levels of protein in all fractions collected using a standard BCA assay. Results are plotted as particles/ml on the left axis and protein (arbitrary units) on the right axis. This demonstrates effectively that the Izon fractions deemed to contain ‘extracellular vesicles’ (fractions 1-4) do not contain significant amounts of protein, which begins to elute from fraction 6 and rises all the way to fraction 12.

**Supplementary Figure 2.**
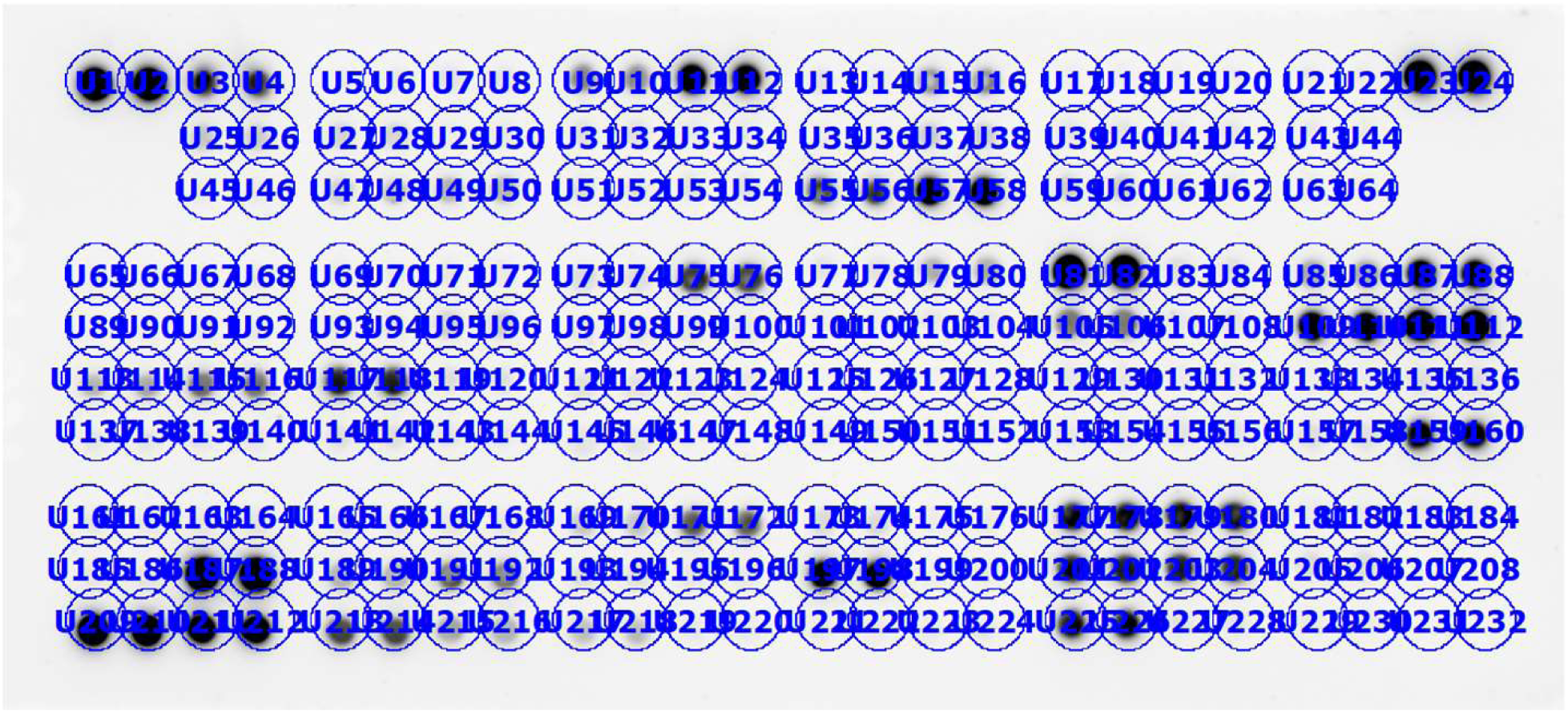
Analysis schematic of the Proteome Profiler. Using Biorad’s ImageLab software, individual circles were drawn initially around the control dots in the top left. These were copied and pasted in order to maintain a consistent size of analysis area and were used to cover all dots on the blot in the pattern outlined by the manufacturer. ImageLab was then used to analyse the amount of staining within each circle.

**Supplementary Figure 3.**
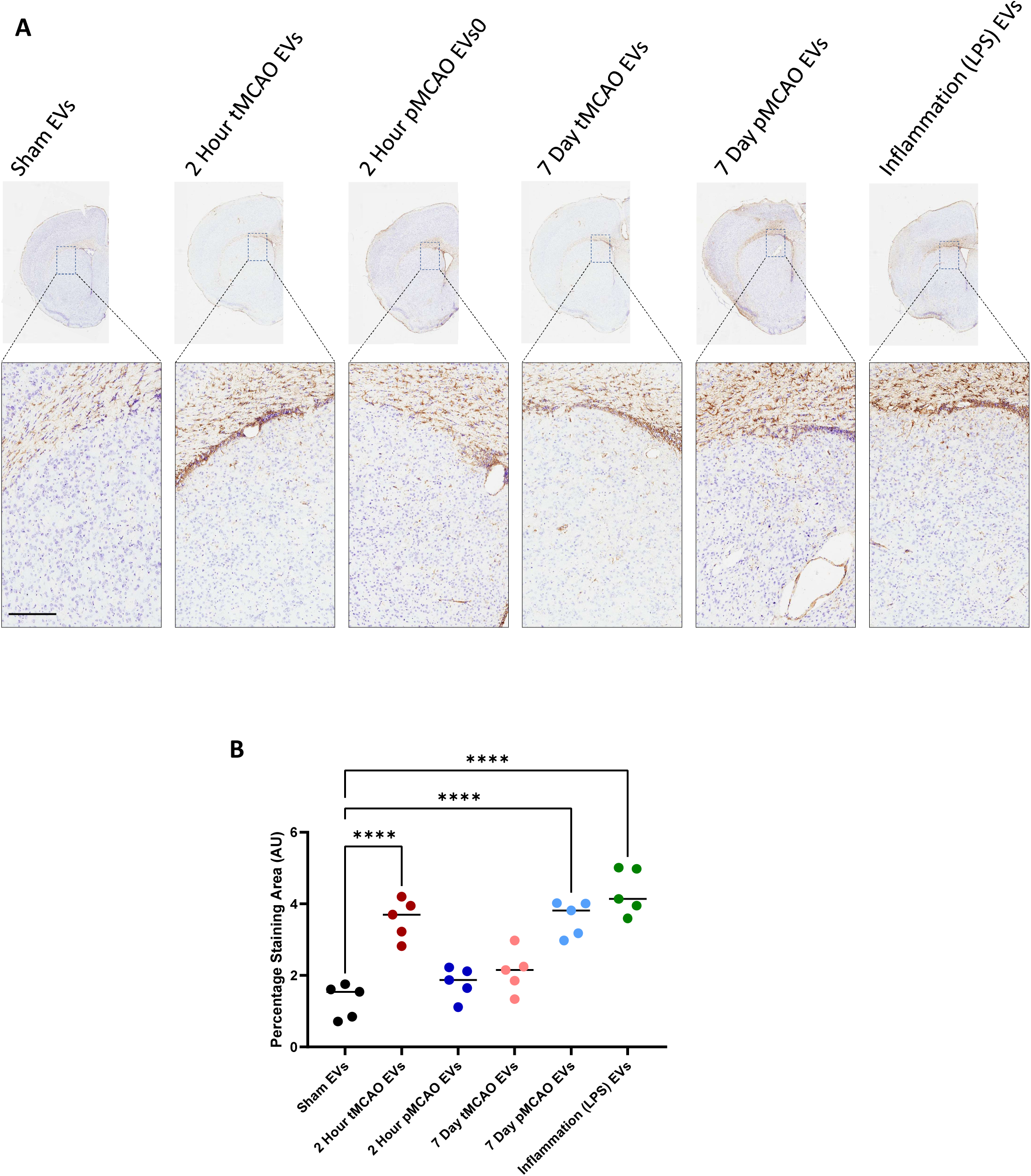
GFAP expression specifically in the corpus callosum after injection of EVs derived from models of stroke and inflammation. After isolation using SEC, EVs from all groups were quantified and 10^10^ EVs from each group were injected in 1µl into the striatum. Tissue was fixed and cryosectioned and stained for GFAP (A). All tissue was also counterstained with cresyl violet. Example images are of the ipsilateral hemisphere (taken from Figure 6) and a magnified area showing detailed staining of GFAP expression within the corpus callosum in each case, individual squares on each image show the area selected for the higher magnification images. Animals injected with EVs from sham animals show very little expression of GFAP, EVs derived from models of stroke and inflammation show varying levels of expression of GFAP (B) with highest corpus callosum expression after injection of EVs from 2 hour tMCAO animals, 7 day pMCAO animals and LPS-challenged animals. Scale bars on images of higher magnification represent 200um. Data are individual animals with mean indicated, n=5. ****p<0.0001.

**Supplementary Figure 4.**
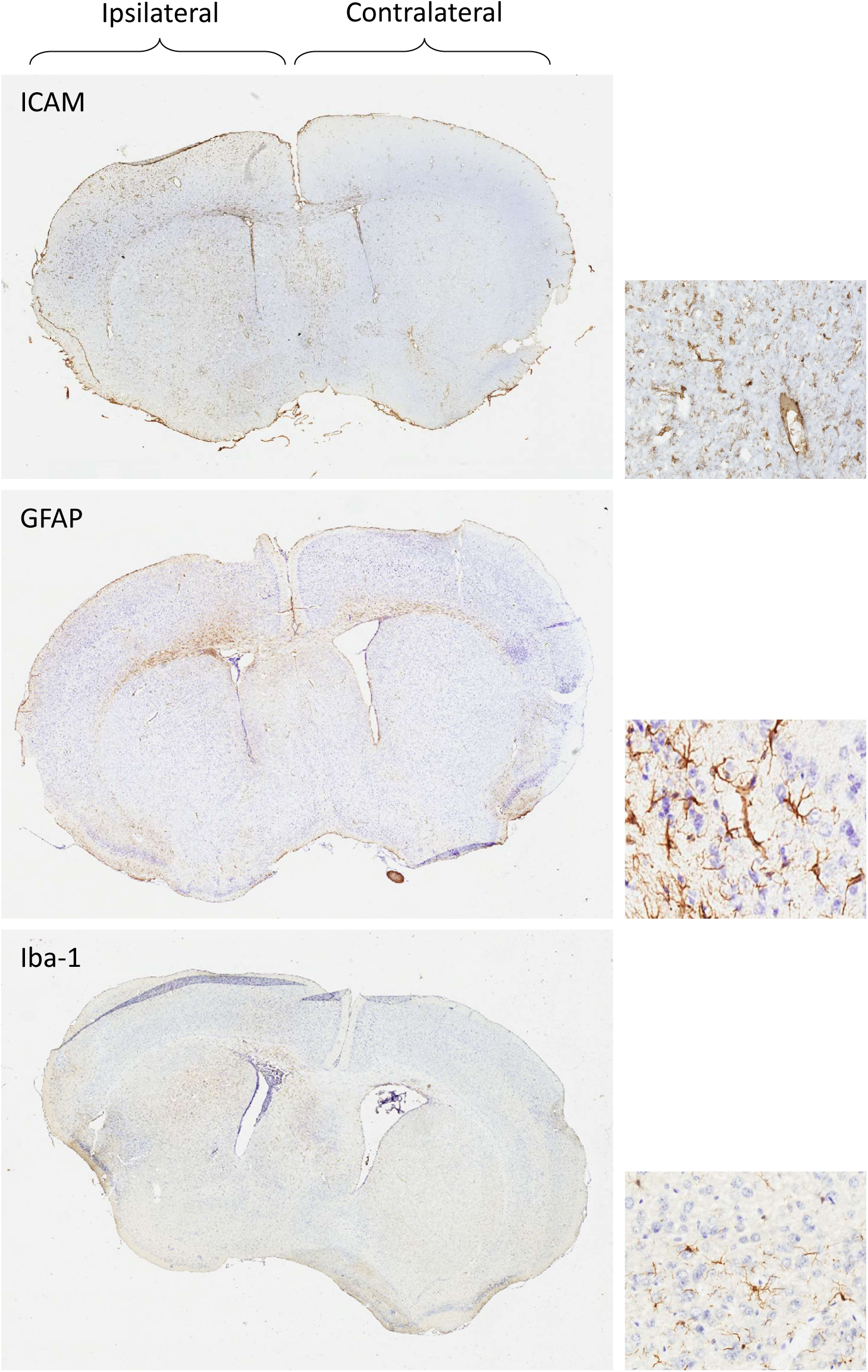
Histological images showing the whole brain to demonstrate the increased staining (particularly of ICAM) in the ipsilateral hemisphere and 20x images to show details of the cells stained. ICAM stains both blood vessels and a variety of glial populations which can be seen in the higher magnification image, GFAP stains astrocytes and the stellate cells are well-represented in the higher magnification image and Iba-1 stains microglia which appear in their amoeboid form in the magnified image.

